# Ageing synaptic vesicles are inactivated by contamination with SNAP25

**DOI:** 10.1101/172239

**Authors:** Sven Truckenbrodt, Abhiyan Viplav, Sebastian Jähne, Angela Vogts, Annette Denker, Hanna Wildhagen, Eugenio F. Fornasiero, Silvio O. Rizzoli

**Affiliations:** Institute for Neuro- and Sensory Physiology, University Medical Center Göttingen, Cluster of Excellence Nanoscale Microscopy and Molecular Physiology of the Brain, Göttingen, Germany; International Max Planck Research School for Molecular Biology, Göttingen, Germany; Master Molecular Biology Programme, University of Vienna, Austria; International Max Planck Research School for Neurosciences, Göttingen, Germany; Leibniz Institute for Baltic Sea Research, Warnemünde, Germany

## Abstract

Old organelles can become a hazard to cellular function, by accumulating molecular damage. Mechanisms that identify aged organelles, and prevent them from participating in cellular reactions, are therefore necessary. We describe here one such mechanism for the synaptic vesicle recycling pathway. Using cultured hippocampal neurons, we found that newly synthesized vesicle proteins were incorporated in the active (recycling) pool, and were preferentially employed in neurotransmitter release. They remained in use for up to ~24 hours, during which they recycled up to a few hundred times. We could only detect one change in the molecular composition of the vesicles, an apparent accumulation of SNAP25 in the aged synaptic vesicles. Overexpression of SNAP25, both in wild-type form or in vesicle-bound form, inhibited exocytosis and promoted the co-localization of the vesicle molecules with a recycling endosome marker. This is in line with the hypothesis that the SNAP25 contamination causes the inactivation of the aged vesicles. The SNAP25 overexpression effect could be alleviated by co-expressing the vesicle-associated molecule CSPα, which has been previously shown to be involved in chaperoning SNAP25 in the vesicle priming process. Overall, these results suggest that newly synthesized vesicle molecules are preferred in vesicle recycling, probably through a mechanism that renders their priming more efficient than that of aged vesicles.

---

Dividing cells are continuously replaced in their entirety, and often remain fully functional even at the end of an organism’s lifespan. In contrast, non-dividing cells, including neurons, accumulate damage to their organelles, which can manifest itself as disease (Sheldrake, 1974; Terman et al., 2007). To prevent detrimental effects, these cells need to avoid the use of aged and damaged organelles, which could disrupt cellular function in unpredictable ways. Such mechanisms have been described, for example, in budding yeast, where old and damaged mitochondria and vacuoles are retained in the ageing mother cell, to prevent their usage in the daughter cells (Nyström and Liu, 2014).

Most mechanisms described so far target damaged components of the organelles, and not their actual age. The damage is dealt with by degrading the organelles, or by sorting out the damaged components. This scenario can function well for cellular processes in which many organelles are used in parallel, and where damage to one organelle is not endangering the outcome of the entire process. It would, however, not be efficient for signaling processes that rely on only a handful of organelles at any one time, such as synaptic vesicle exocytosis (Denker et al., 2011a; Harata et al., 2001; de Lange et al., 2003; Richards et al., 2003). Such processes could be severely disrupted by single damaged vesicles. If one or more of the synaptic vesicles docked at an active zone fail to release, the synapse remains silent, and fails to transmit the signal. Even more importantly, if released vesicles fail to recycle correctly, they might fail to liberate the active zone, which would result in a persistent inhibition of the synapse (Haucke et al., 2011). This suggests that neurons should have mechanisms in place that recognize old, damage-prone vesicles, to inhibit the use of such vesicles in synaptic release, even before they accumulate substantial levels of damage.

Some evidence in support of this hypothesis exists in non-neuronal secretory cells, including chromaffin cells (Duncan et al., 2003) and pancreatic β-cells (Ivanova et al., 2013). Here, newly synthesized dense-core vesicles, which are not recycled after use, appear to be more release-prone than aged ones, since the former are released during mild (presumably more physiological) cellular activity, while the latter can only be exocytosed after heavy artificial stimulation. The situation affecting synaptic vesicles, which recycle repeatedly within the synaptic boutons, is less clear. Here, the vesicles can be broadly separated into 1) an active recycling pool, which includes the readily releasable vesicles that are docked at the release sites, and 2) an inactive reserve pool that participates little in release under most stimulation conditions, and can typically only be released by heavy action potential stimulation (Rizzoli and Betz, 2005), by pharmacological manipulations (Kim and Ryan, 2010), and/or by the use of drugs that increase substantially presynaptic activity (Frontali et al., 1976).

It is currently unclear whether the situation observed with non-recycling dense-core vesicles mirrors the situation from synaptic boutons, with the reserve pool representing an aged population. However, a further issue complicates such an interpretation: the problem of the vesicle identity (Ceccarelli and Hurlbut, 1980; Haucke et al., 2011; Rizzoli, 2014), for which two opposing models have been presented. In a first model, the vesicle maintains its protein composition after exocytosis, as a single patch of molecules on the plasma membrane, which is then retrieved as a whole by endocytosis. In the second model, the vesicle loses its molecular cohesion upon fusion, and its proteins diffuse in the plasma membrane and intermix with other vesicle proteins, before endocytosis. In the first scenario, the neuron could readily target old vesicles for removal. This is less obvious in the second scenario, since vesicles lose their identity by intermixing molecules, which makes it more difficult to pinpoint the old vesicle components.

Although it has been difficult to differentiate between these two scenarios, an unified view is starting to emerge. This view suggests that several synaptic vesicle proteins remain together during recycling, as meta-stable molecular assemblies, although not as whole individual vesicles (Rizzoli, 2014). In simple terms, this view implies that the vesicle splits into a number of meta-stable protein assemblies after exocytosis. These individual assemblies are stabilized by strong interactions between abundant vesicle proteins such as synaptophysin and synaptobrevin/VAMP2 (Adams et al., 2015; Becher et al., 1999; Mitter et al., 2003; Pennuto et al., 2003), are strong enough to persist even after detergent solubilization (Bennett et al., 1992), and may be further stabilized by an interaction with the endocytosis machinery (Gimber et al., 2015). The assemblies have been observed by all laboratories that have studied this issue using super-resolution imaging (see for example Hoopmann et al., 2010; Hua et al., 2011; Opazo et al., 2010; Wienisch and Klingauf, 2006; Willig et al., 2006), and are fully compatible with modern interpretations on the meta-stable nature of membrane protein assemblies (for example Saka et al., 2014a). The vesicle protein assemblies are then regrouped during endocytosis and vesicle reformation, resulting in new synaptic vesicles.

Could such a scenario enable a neuron to nevertheless distinguish between old and young vesicles? This seems unlikely at the level of the single vesicles, but entirely possible at the level of the vesicle pools. The active and inactive synaptic vesicles maintain their pool identities over long time periods (see review Rizzoli and Betz, 2005), and it has been thoroughly demonstrated that mild (physiological) stimulation results in no molecular mixing among vesicles from different pools (Wienisch and Klingauf, 2006). Therefore, as long as the recycling vesicles only intermix molecules with other recycling vesicles, they remain separated from the reserve ones, and could be specifically targeted and/or timed by the neurons. At the same time, the meta-stable nature of the vesicle protein assemblies implies that such assemblies could become labeled over time with markers that could tag them as old, used molecular patches (markers such as non-vesicle molecules or damaged, unfolded proteins).

The hypothesis that neurons have mechanisms to recognize old and damage-prone vesicles is thus plausible, irrespective of whether the vesicle fully maintains its identity during recycling, or splits into meta-stable assemblies that intermix within one single pool of vesicles (within the recycling pool). We set out to test this hypothesis here, and found that ageing recycling vesicle proteins were indeed identified by the neurons. We used multiple assays to test the hypothesis that the recycling vesicle proteins are metabolically younger than the non-recycling reserve vesicles. This was indeed what we observed in all assays we used. We also found that non-recycling, aged vesicles co-localized more strongly with SNAP25, a plasma membrane molecule that is typically not found in vesicles at high levels (Takamori et al., 2006). SNAP25 expression reduced exocytosis, which suggests that SNAP25 may be able to render the aged vesicles less prone to release. As this effect could be removed by expressing the priming chaperone CSPα, a simple hypothesis is that the SNAP25 accumulation reduces the aged vesicle exocytosis by interfering with the priming step performed by CSPα.

## RESULTS

### Synaptic vesicles protein assemblies on the plasma membrane

As mentioned in the Introduction, several laboratories suggested that synaptic vesicle proteins form meta-stable clusters on the plasma membrane after exocytosis, which are then taken up by endocytosis, and therefore remain together, as groups of molecules, throughout synaptic vesicle recycling. This has been the case especially for limited stimulation levels that are comparable to the intrinsic network activity of the cultures (Fernández-Alfonso et al., 2006; Hoopmann et al., 2010; Hua et al., 2011; Opazo et al., 2010; Wienisch and Klingauf, 2006; Willig et al., 2006).

Nevertheless, we set out to test this again, before proceeding to further experiments, since this is a critical issue for the reasoning presented in the Introduction. We used hippocampal neurons that were mildly stimulated, by depolarizing for 6 minutes with 15 mM KCl, in presence of fluorophore-conjugated antibodies directed against the lumenal domains of the vesicle proteins synaptotagmin 1 (Kraszewski et al., 1995; Matteoli et al., 1992), which reveal synaptotagmin 1 molecules present on the plasma membrane. To ensure that the antibodies only revealed synaptotagmin 1 molecules that were exocytosed during stimulation, and not epitopes that were already present on the plasma membrane, we blocked these epitopes by incubating the neurons with non-fluorophore-conjugated antibodies before application of the stimulation pulse and the fluorophore-conjugated antibodies (Hoopmann et al., 2010). The fluorophore-conjugated antibodies could thus only bind to the synaptotagmin molecules from newly exocytosed vesicles. To determine whether such molecules diffused away from each other, and/or from other vesicle proteins, we fixed the neurons and immunostained for another synaptic vesicle marker, synaptophysin, using an antibody that recognized the lumenal domain of synaptophysin (Hoopmann et al., 2010). The immunostaining was performed without permeabilization, so that only surface synaptophysin molecules could be detected.

We then quantified the co-localization of synaptotagmin 1 and synaptophysin on the membrane surface by two-color super-resolution STED microscopy (Hoopmann et al., 2010; Supplementary Fig. 1a). We found that the majority of synaptotagmin 1 and synaptophysin signals co-localized, indicating that the molecules are found in synaptic vesicle protein assemblies or co-clusters. Only a small proportion of the synaptotagmin 1 signals was found away from the synaptophysin signals, and thus may have corresponded to molecules lost from the synaptic vesicle protein assemblies. As discussed in the Introduction, we expected that assemblies of vesicle molecules, containing different types of vesicle proteins, would appear on the plasma membrane after exocytosis, and that only a minority of the proteins would diffuse away and be lost in the membrane, far from the assemblies. To test this, we tagged the synaptic vesicle proteins synaptophysin and synaptotagmin 1 from newly exocytosed vesicles, using fluorescently conjugated antibodies, and quantified their co-clustering We observed a very limited loss of proteins from the synaptic vesicle protein assemblies (co-clusters), (around 3%, Supplementary Fig. 1b,c). We therefore conclude, in line with the previous literature, that newly exocytosed synaptic vesicle proteins appear to form co-clustered assemblies on the plasma membrane.

### Synaptic vesicles become inactive as they age

To follow synaptic vesicle proteins through their life cycle in the synapse, we reasoned that a similar use of antibodies directed against the lumenal domain of synaptic vesicle proteins could reveal the localization and activity status of such vesicle molecules for substantial time periods. We started with the same fluorophore-conjugated synaptotagmin 1 antibodies as above. This procedure is highly specific, as antibodies not directed against surface proteins of the neurons are not taken up efficiently by the neurons, resulting in essentially null backgrounds (Supplementary Fig. 2a). At the same time, the synaptotagmin 1 604.2 antibody we used here binds its epitope with remarkable stability. The antibodies remained bound onto fixed neurons for up to 10 days, with no noticeable loss of signal, even when incubated with a 100x molar excess of antigenic peptide (Supplementary Fig. 2b). This large excess ensures that antibodies that come off their targets would immediately bind the peptides, and would not be able to find the cellular targets again. This did not happen, even when the fixed neurons were incubated at pH 5.5 during the entire procedure (the pH encountered by the antibodies within vesicles). This implies that these antibodies bind stably to their targets, and are not affected by the vesicular pH. At the same time, these antibodies (this clone, 604.2) have been used in multiple experiments in synaptic vesicle recycling, and appear not to interfere with it (Hoopmann et al., 2010; Hua et al., 2011; Opazo et al., 2010; Wienisch and Klingauf, 2006; Willig et al., 2006). We also observed the tagged synaptic vesicles to be fully responsive to stimulation (Supplementary Fig. 2c), demonstrating that these antibodies were taken up specifically by synaptic vesicles in the conditions of labeling that we used (1 hour of antibody incubation during intrinsic network activity).

To check the behavior of the antibody-labeled molecules, we applied the synaptotagmin 1 antibodies on living, active rat primary hippocampal cultures (Fig. 1a). The antibodies, which were conjugated to the bright, pH-insensitive fluorophore Atto647N, were taken up during the intrinsic network activity of the cultures (Fig. 1b) and co-localized well with synaptic vesicles for up to 10 days (Supplementary Fig. 3a,b). Incubating the neurons with the antibodies for one hour was sufficient to tag most of the epitopes in the actively recycling pool of vesicles (Supplementary Fig. 3c), as further incubation beyond 30 minutes did not result in significant additional labeling (Supplementary Fig. 2d). To quantify the proportion of the synaptotagmin 1 molecules that were labeled by this procedure, we compared the resulting images with those of neuronal cultures that were immunostained for synaptotagmin 1 with the same antibody, after fixation and permeabilization, and in which all epitopes were thus revealed. This showed that the live labeling procedure revealed approximately 50% of the synaptotagmin 1 epitopes (Supplementary Fig. 3c). This is in agreement with previous findings on the size of the recycling pool in these cultures (Rizzoli and Betz, 2005). Approximately half of the labeled epitopes were found on the surface membrane, as part of the readily retrievable pool, again in agreement with previous findings (Wienisch and Klingauf, 2006).

**Figure 1:**
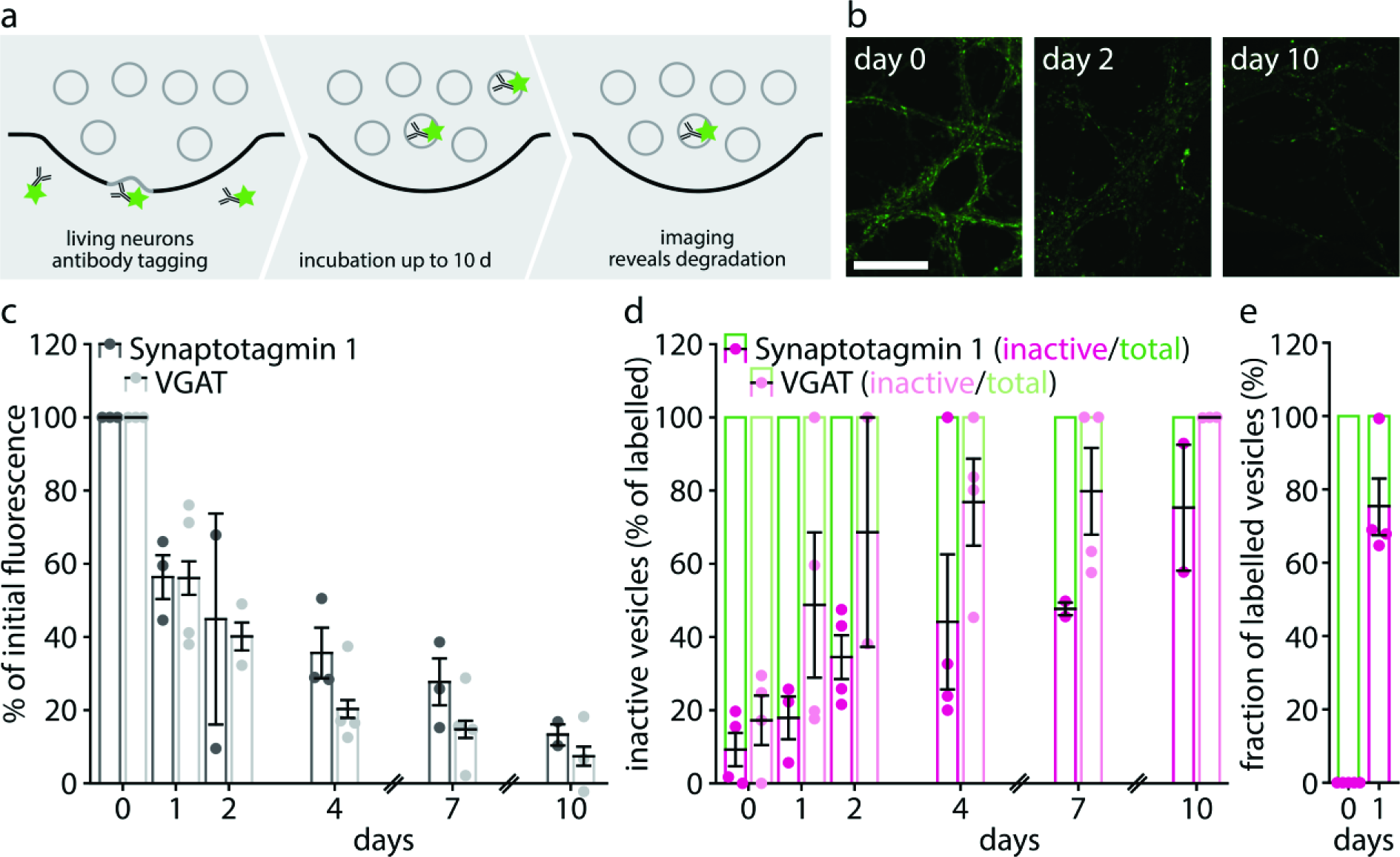
Ageing synaptic vesicle proteins stop releasing. (a) To determine the time interval that synaptic vesicle proteins spend in synapses, we incubated living hippocampal cultured neurons with fluorophore-conjugated (green stars) antibodies against the lumenal domain of synaptic vesicle proteins. Fluorophore-conjugated synaptotagmin 1 (Atto647N-conjugated) or VGAT (CypHer5E-conjugated) antibodies were applied (diluted 1:120 from 1 mg/ml stock) to the neurons in their own culture medium, for 1 h at 37°C in a cell culture incubator. This interval is sufficient to label the molecules from the pool of recycling vesicles (Supplementary Figure 2c). The antibodies were then washed off, and the neurons were placed again in the incubator. Individual coverslips were then retrieved at different time intervals after labeling, fixed, and imaged, to determine the amount of synaptotagmin 1 and VGAT labeling remaining in synapses. (b) Exemplary images of neurons labelled with synaptotagmin 1 antibodies, imaged at different time points after labeling. Scale bar: 20 μm. (c) The loss of synaptic vesicle proteins from the synapse was monitored by imaging the synaptotagmin 1 or VGAT antibody fluorescence at serial time points after tagging synaptotagmin 1 (dark grey; n = 3, 3, 2, 3, 3, 2 independent experiments per respective time point, at least 10 neurons sampled per experiment) or VGAT (light grey; n = 3, 4, 2, 4, 4, 3 independent experiments per respective time point, at least 10 neurons sampled per experiment). To determine fluorescence intensities specifically in synapses, the neurons were immunostained for the synaptic vesicle marker synaptophysin, and only the fluorescence found within synaptophysin spots (synapses) was analyzed. We did not observe any significant changes in the co-localization of fluorophore-conjugated antibodies in respect to synaptophysin over time (Supplementary Fig. 2a,b). (d) To determine whether the antibody-tagged synaptic vesicle proteins that are present in synapses at different time intervals after labelling are exocytosis-competent, we labeled neurons with CypHer5E-conjugated antibodies for synaptotagmin 1 or VGAT, exactly as described in (a). At different time intervals after labeling, individual cultures were removed from the incubator, mounted in a live-stimulation chamber, and subjected to a stimulation paradigm designed to release the entire recycling pool (20 Hz, 30 s; Wilhelm et al., 2010). Exocytosis is determined by monitoring the fluorescence of the pH-sensitive fluorophore CypHer5E, which is fluorescent within vesicles, but is quenched by exposure to the neutral extracellular pH. To determine the proportion of the labeled vesicles that is still able to exocytose, we measured the fluorescence signal corresponding to all labeled vesicles by fixing and permeabilizing the neurons, and then exposing them to a pH of 5.5, obtained with TES buffered solutions. The following number of experiments were performed: for synaptotagmin 1 (dark green and dark magenta) n = 4, 3, 4, 4, 2, 2 independent experiments per respective time point, with at least 12 neurons sampled per experiment); for VGAT (light green and light magenta) n = 4, 4, 2, 4, 4, 3 independent experiments per respective time point, with at least 8 neurons sampled per experiment). The VGAT signal follows the same approximate dynamic as synaptotagmin 1, but is faster. VGAT is present in only ~5-10% of the neurons in our cultures, and therefore this difference may reflect a cell type-specific effect. (e) A similar experiment, based on synaptotagmin 1 antibodies, was used to determine the fraction of vesicles of different ages that recycled during intrinsic network activity, in the absence of external stimulation. This experiment is shown fully in Supplementary Fig. 3, where the procedure is described in detail. Synaptotagmin 1 antibodies were preferred, since only a minority of the neurons in our cultures are VGAT-positive, and therefore they are not representative for the general spontaneous network activity of the cultures. The fraction of vesicle proteins that are still releasable under intrinsic network activity is only about half of that which can respond to high-frequency stimulation after 1 day. (n = 5 and 4 independent experiments per respective time point, at least 10 neurons sampled per experiment). All data represent the mean ± SEM.

Having verified that the synaptotagmin 1 antibodies label a pool of ~50% of the synaptic molecules, which are distributed both on the surface (as readily retrievable molecules) and within the bouton (as readily releasable vesicles), we next investigated the behavior of the labeled molecules over time. We labeled a series of neuronal cultures at one time point, and then removed coverslips for fixation after different time intervals, of up to 10 days. We then immunostained the neurons for synaptophysin, an optimal marker for synaptic vesicles (and thereby synaptic boutons), and analyzed the intensity of the synaptotagmin 1 antibody signals within the synaptophysin-marked boutons. We found that the synaptotagmin 1 signals decreased slowly (Fig. 1c). Within ~2 days the synaptotagmin 1 signals could be seen in large acidified compartments in the cell bodies, (Supplementary Fig. 3d). This suggests that synaptotagmin 1 molecules were degraded through a lysosomal route, in agreement with the literature (Rizzoli, 2014). This suggestion was further confirmed by the observation that inhibiting lysosomal degradation using leupeptin resulted in a decrease in the loss of synaptotagmin 1 from boutons (Supplementary Fig. 3e). The rate of the synaptotagmin 1 loss was well within the range previously measured for synaptic vesicle proteins by radioisotopic labeling or mass spectrometry (Cohen et al., 2013; Daly and Ziff, 1997).

To test whether this phenomenon was also observable for other vesicle molecules, we turned to antibodies against the vesicular GABA transporter, VGAT (Fig. 1c). This is the only other vesicle protein for which lumenal antibodies are available (Martens et al., 2008), as the synaptophysin antibodies used in Supplementary Fig. 1 do not recognize the unfixed protein, and cannot be used in live labeling. The loss of VGAT antibodies from synapses paralleled that of synaptotagmin antibodies (Fig. 1c).

To test whether the tagged proteins of different ages were still involved in synaptic activity, we employed antibodies conjugated to the pH-sensitive dye CypHer5E (Hua et al., 2011; Martens et al., 2008). We used a stimulation protocol designed to trigger the release of the entire population of recycling vesicles, 600 action potentials at 20 Hz (Wilhelm et al., 2010), and we measured the fraction of the CypHer5E-labeled proteins that were releasable (Fig. 1d). This was monitored through imaging the stimulation-induced reduction in CypHer5E fluorescence, which is quenched at neutral pH, and therefore becomes invisible upon exocytosis. We found that the fraction of the labelled proteins that could be induced to release gradually decreased with age, until almost no response could be elicited any more, at 7-10 days after tagging (Fig. 1d). This observation was made for both synaptotagmin 1 and VGAT. A trivial explanation for this observation would be that the neurons became slowly damaged during the 7-10 days, simply by ageing, resulting in decreased release activity. This, however, was not the case, as the neurons were still active throughout the entire study period, and recycled constant amounts of vesicles (Supplementary Fig. 2e).

We also investigated this phenomenon during intrinsic network activity, focusing on synaptotagmin 1 (since the low proportion of VGAT-positive neurons in the culture, ~5-10%, is likely not representative for the general activity of our cultures). We performed here a different experiment (Supplementary Fig. 3a), as continuous imaging for long time periods, as required for the limited normal activity of the cultures, is not possible with CypHer5E, which is readily bleached. We tagged synaptotagmin 1 molecules with non-fluorescently conjugated antibodies at one time point, and then, after different time intervals, of up to several days, we applied green-conjugated secondary antibodies onto the living cultures. These antibodies reveal all of the synaptotagmin 1 antibodies that are exposed during vesicle recycling. After incubating the living cultures with the secondary antibodies for one hour, we fixed and permeabilized them, and applied red-conjugated secondary antibodies, to thereby reveal all of the remaining synaptotagmin 1 antibodies. This procedure thus indicates the proportion of the tagged synaptotagmin 1 molecules that are still recycling in the synapses under normal activity (in green), as well as the proportion that are present in the synapses (in magenta). The proportion of the molecules that participated in release decreased relatively fast, within about one day (time constant of ~0.4 days; Supplementary Fig. 3b,c). The molecules were not lost as rapidly from the synapses, and were found there for substantially longer (Supplementary Fig. 3b,c).

Thus, we concluded that synaptotagmin 1 molecules became rapidly inactive (from the point of view of exocytosis), albeit many still remained within synapses. Such vesicles could be triggered to release by strong stimulation (Fig. 1d), but did not release under normal network activity (Fig. 1e, Supplementary Fig. 3b,c). For simplicity, we termed this the “inactive state” of synaptic vesicles, or vesicle molecules. It could also be called a “reserve” or “reluctant” state: the vesicles are present in the synapse, release upon supra-physiological stimulation, but not during normal activity. The inactivity (or reluctance) became absolute after about 7-10 days (Fig. 1d).

### A sensor for the age of vesicle proteins confirms that young molecules are preferentially employed in exocytosis

To complement these antibody-based approaches, we tested the behavior of the synaptic vesicle protein VAMP2 after tagging with a novel construct that enables the separate identification of newly synthesized or older proteins. We expressed VAMP2 coupled to a SNAP tag (Keppler et al., 2003, 2004), and separated from the original sequence by a TEV protease cleavage site: VAMP2-TEV-SNAPtag (Supplementary Fig. 4a). This construct should be minimally disruptive to physiological synaptic vesicle function, since VAMP2 is by far the most abundant synaptic vesicle protein (Takamori et al., 2006; Wilhelm et al., 2014), and therefore every vesicle should still have ample levels of wild-type VAMP2, independent of the levels of VAMP2-TEV-SNAPtag expression. Moreover, our construct is designed on the basis of synaptopHluorin (VAMP2-pHluorin), which is known to target and function well in neurons (Gandhi and Stevens, 2003; Miesenböck et al., 1998).

The construct is then expressed, ensuring that synaptic vesicles are tagged with it, but it is not inherently fluorescent. When desired, the construct can be revealed by labeling with a membrane-permeable fluorophore, such as tetramethyl-rhodamine-Star (TMR-Star). The coupling reaction is self-catalyzed by the SNAP tag (Juillerat et al., 2003), and is highly efficient, as the application of a second SNAP-binding fluorophore immediately afterwards results in no detectable signal. This, therefore, labels all SNAP-tagged proteins present at a particular time point in the cells. After allowing protein biosynthesis to proceed for a further 1-2 days, the newly produced, younger constructs can be revealed by labeling with a second fluorophore, such as the 647-SiR, which is easily spectrally separable from TMR-Star. This thus reveals the younger VAMP2 molecules. The neurons now contain two populations of VAMP2-TEV-SNAPtag, one young, labeled with 647-SiR, and one that is 1-2 days older, labeled with TMR-Star (Supplementary Fig. 4b). To test whether young or old vesicles are preferentially used in recycling, one can incubate the neurons with TEV protease, which cleaves the fluorescent tags away from vesicles that participate in recycling. Inactive vesicles are not affected by the TEV protease, since it cannot penetrate the cell membrane. Upon adding the protease, the 647-SiR signals were preferentially reduced, indicating that the tag was cleaved preferentially from young VAMP2 proteins (Supplementary Fig. 4b,c).

### Releasable synaptic vesicles are metabolically younger than inactive vesicles

Having verified with two independent techniques that newly synthesized protein copies are preferentially used in exocytosis for three vesicle proteins (synaptotagmin 1, VGAT, VAMP2), we proceeded to test whether the entire protein makeup of actively recycling vesicles is metabolically younger than that of non-recycling, inactive vesicles. This can only be performed by imaging the makeup of the vesicles, after tagging all proteins with a marker for their turnover. This type of procedure can be performed by incubating the cultures with a special amino acid, which is incorporated in all of the newly biosynthesized proteins, and serves therefore as a marker for newly made proteins. Two techniques are available for this type of tagging. First, cells can be fed with an unnatural amino acid, such as azidohomoalanine (AHA), which incorporates into newly produced proteins in the position of methionine, and can be detected in fluorescence microscopy after fluorophore conjugation (FUNCAT; Dieterich et al., 2011), through a highly specific azid-alkyne reaction that has been termed CLICK chemistry (Rostovtsev et al., 2002). Second, the cells can be fed with an amino acid, such as leucine, that contains a stable rare isotope, such as ^15^N. The isotope can be detected through the mass spectrometry imaging technique nanoSIMS (Lechene et al., 2006; Steinhauser and Lechene, 2013), which has a higher resolution than conventional fluorescence microscopy (~50-100 nm in cultured neurons, Saka et al., 2014b).

We applied here both techniques, in separate experiments, tagging newly synthesized proteins either with AHA or with leucine containing ^15^N. We then incubated the cells with synaptotagmin 1 antibodies, to label the recycling vesicles. Alternatively, we incubated the neurons with the antibodies several days before feeding them with the special amino acids, to make sure that the tagged vesicles were already in the inactive state at the time of amino acid incorporation into newly synthesized proteins. We then imaged the vesicles in fluorescence microscopy, and correlated their positions with those of the AHA or ^15^N signals, obtained in STED microscopy or in nanoSIMS, respectively (Saka et al., 2014b).

In both approaches, we could detect a significantly higher co-localization of the actively recycling synaptic vesicles with newly synthesized proteins (Fig. 2b,c). Moreover, the higher sensitivity of nanoSIMS, which, unlike FUNCAT, detects simultaneously both the ^14^N from old proteins and the ^15^N from new proteins, could demonstrate that the active vesicles were significantly younger than the rest of the axon. The opposite was true for the inactive vesicles (Fig. 2c).

**Figure 2:**
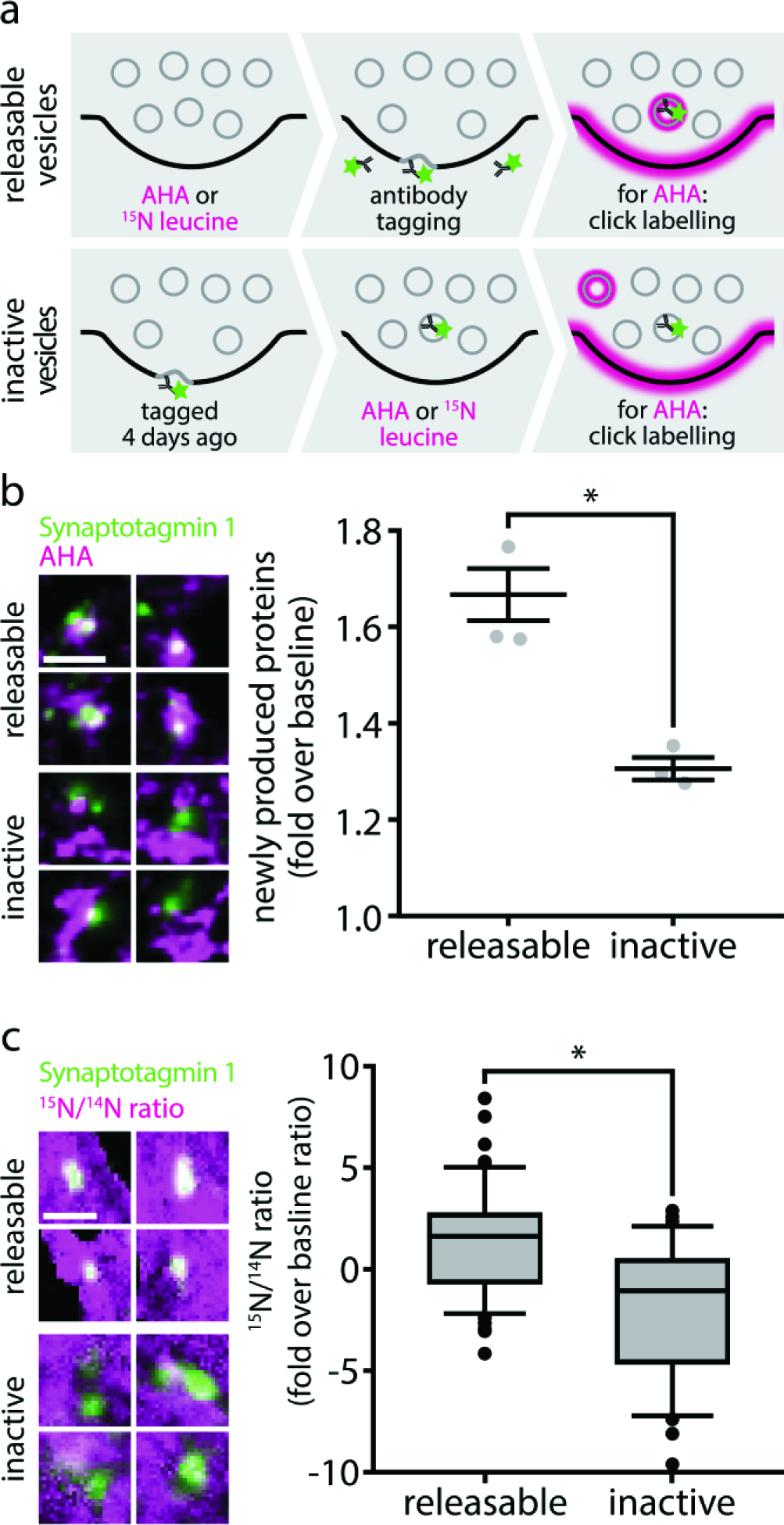
Metabolic imaging reveals that releasable synaptic vesicles are younger than inactive vesicles. (a) To label newly produced proteins, we used the unnatural amino acid AHA, which was detected in STED imaging, or ^15^N-leucine, which was detected in nanoSIMS imaging. We used two experimental paradigms for this. First, to correlate the presence of newly synthesized proteins with recycling vesicles, we fed these compounds to neurons for 9 hours (AHA) or 24-72 hours (^15^N-leucine), before labeling the proteins from the recycling pool by applying synaptotagmin 1 antibodies to the neurons, for 1 hour at 37°C, as in Figure 1a. AHA was applied in culture medium free of methionine, which is the amino acid AHA replaces during protein biogenesis. ^15^N leucine was fed to the cultured neurons in 3-fold molar excess over ^14^N leucine in their normal culture medium for 24-72 hours prior to processing the samples. The extended feeding time compared to AHA was necessary to reliably obtain a signal in nanoSIMS imaging. Second, to compare the co-localization of newly synthesized proteins with inactive synaptic vesicles, we performed the same experiment but with 3-4 days between antibody tagging and metabolic labeling. We labeled the recycling vesicles by synaptotagmin 1 antibody incubation, as in Figure 1a, and then returned the cell cultures to the incubator. We then waited for 4 days for the inactivation of the vesicles to take place, and then fed the neurons with AHA or ^15^N, to label newly synthesized proteins. (b) Using this approach, we analyzed the co-localization of AHA with releasable or inactive synaptic vesicles in STED microscopy. The neurons were fixed and subjected to a click chemistry procedure that reveals all AHA moieties, coupling them to the fluorophore Chromeo494 (a procedure known as FUNCAT; Cohen et al., 2013). The neurons were then embedded in melamine resin, and were sectioned into 20 nm sections on an ultramicrotome. The sections were then imaged by 2-color STED microscopy. The amount of AHA fluorescence co-localizing with the releasable or inactve vesicles was determined, and was expressed as fold over baseline signals. n = 3 independent experiments per data point, at least 10 neurons sampled per experiment, *p = 0.0037; t(4) = 6.0980). Scale bar: 500 nm. (c) We further analyzed the co-localization of AHA with releasable and inactive synaptic vesicles in correlated fluorescence and isotopic microscopy (COIN), as follows. The neurons were fixed and were embedded in LR White resin, a vinyl resin that is usable in both fluorescence and isotopic secondary ion mass spectrometry imaging. The samples were then sectioned into 200 nm sections on an ultramicrotome, as this thickness is ideal for secondary ion mass spectrometry. The sections were mounted on silicon wafers, and were imaged on a Nikon Ti-E microscope, using 150x magnification, to detect the synaptic vesicles. The same areas were then imaged in a nanoSIMS instrument, recording both the ^15^N and ^14^N signals. The ^15^N/^14^N ratio was then determined both within the vesicle areas and elsewhere, and was then presented as fold over the baseline ratio in the axons. n = 57 synapses from 3 independent experiments for releasable synaptic vesicles, n = 47 synapses from 2 independent experiments for inactive vesicles, *p = 0.0001, t(102) = 5.5378. The ratio is a direct indication of the amount of ^15^N-leucine in the vesicles. The inactive vesicles contain substantially fewer newly synthesized proteins than the rest of the axon (the ^15^N/^14^N ratio of which served as baseline; p = 0.0001, t(96) = 4.0691), while the releasable ones contain substantially more newly synthesized proteins (p = 0.0004, t(116) = 3.6156). Scale bar: 1 μm. All data represent the mean ± SEM.

### Inactivated vesicles cannot replace young vesicles in the releasable population

We then tested whether synaptic vesicles, once inactivated, could return to the recycling pool under normal culture activity. Strong stimulation, as in Fig. 1d, enables such vesicles to release, albeit they appeared not to do it under normal activity (Supplementary Fig. 2). To test this, we applied unconjugated lumenal synaptotagmin 1 antibodies to saturate all of the epitopes of the releasable population (Fig. 3a), and then followed this up with pulses of fluorophore-conjugated synaptotagmin 1 antibody, to reveal new epitopes entering the active, releasable population (Fig. 3b,c).

**Figure 3:**
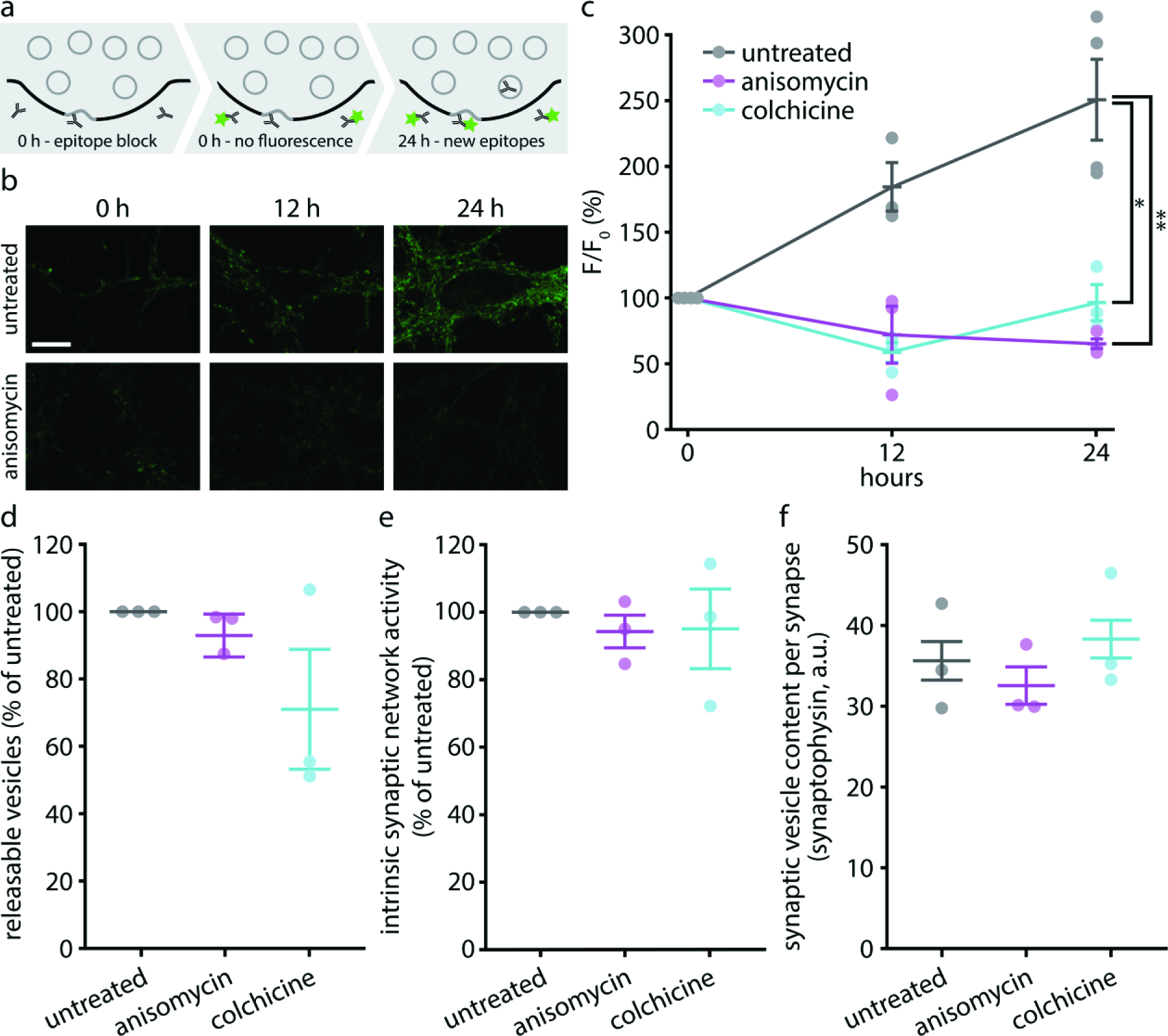
The synapse is dependent on young synaptic vesicles to turn over the active population. (a) To determine the rate at which new synaptotagmin 1 epitopes come into the recycling pool, we saturated the lumenal synaptotagmin 1 epitopes on active synaptic vesicles by incubating with an unconjugated monoclonal antibody for 2 hours at 37°C, and then followed the appearance of new epitopes by applying a fluorophore-conjugated (Atto647N) version of the same monoclonal antibody, at different time points after saturating the initial epitopes. Individual coverslips were only used to investigate single time points, being fixed before imaging. (b) Exemplary images taken after applying the fluorophore-conjugated antibodies at 0, 12 or 24 hours after saturating the initial synaptotagmin 1 epitopes. Scale bar: 10 μm. The images were taken with a Leica confocal microscope. (c) The same experiment was performed, but the cultures were incubated with drugs that disrupt protein synthesis (anisomycin) or microtubule-based transport (colchicine). The fluorescence was measured at different time points, and expressed in % of the initial fluorescence at the initial time point. n = 4 independent experiments per data point for untreated 0 h and 24 h, n = 3 for all else, at least 10 neurons sampled per experiment; *p = 0.0104, t(5) = 3.9882; **p = 0.0041, t(5) = 5.0091. (d) To determine whether the drugs impair synaptic activity, neurons were treated with the drugs for 24 h, and were then stimulated in the presence of fluorescently conjugated synaptotagmin 1 antibodies (20 Hz, 30 s), to label the entire recycling pool. Only a limited decrease in recycling was observed in drug-treated preparations. (n = 3 independent experiments per data point, at least 10 neurons sampled per experiment; untreated vs. anisomycin, p = 0.2048, t(4) = 1.5131; untreated vs. colchicine, p = 0.1787, t(4) = 1.6286). (e) Drug effects on the intrinsic network activity. To assess the intrinsic network activity of our cultures after 24 h of drug treatment, we measured vesicle recycling during the last hour of the treatment. The network activity did not change significantly (n = 3 independent experiments per data point, at least 10 neurons sampled per experiment; untreated vs. anisomycin, p = 0.3317, t(4) = 1.1035; untreated vs. colchicine, p = 0.6998, t(4) = 0.4144). (f) The total synaptic vesicle pool size after 24 h of drug treatment. This value was determined by immunostaining for a major synaptic vesicle marker, synaptophysin. There were no significant changes in synaptic vesicle pool size (n = 3 independent experiments per data point, at least 10 neurons sampled per experiment; untreated vs. anisomycin, p = 0.5374, t(4) = 0.6738; untreated vs. colchicine, p = 0.6566, t(4) = 0.4796). All data represent the mean ± SEM. Imaging was performed with a Leica confocal microscope.

These vesicles could come from two sources: newly synthesized vesicles from the cell body, or the inactivated vesicles, whose epitopes are not affected by the initial incubation with unconjugated antibodies, and which account for ~50% of all of the vesicles in the synapse (Supplementary Fig. 2c). Cutting off the production of new synaptic vesicles by blocking protein biosynthesis with anisomycin, or by disrupting the microtubule transport network with colchicine, completely removed the entry of new epitopes into the releasable population (Fig. 3b,c). These drugs were applied after the tagging with unconjugated antibodies and remained present until application of the conjugated antibody.

This indicates that, under conditions of intrinsic network activity, the synapse is dependent on young vesicles to replace its releasable population. The drugs did not significantly affect the proportion of releasable vesicles in the synapse, the intrinsic network activity, or the total amount of vesicles per bouton (Fig. 3d-f), suggesting that the neurons were still healthy at the time of the experiments.

### Increased synaptic activity accelerates ageing and inactivation

We next investigated whether temporal age is the defining factor for inactivating vesicles, or whether the usage of the vesicles (or vesicle proteins) is responsible, and, if the latter is true, how often proteins from the vesicles could be used before inactivation.

If the age of the vesicle proteins is the defining parameter in the inactivation, then increasing the frequency of vesicle release and recycling should have no influence on the rate of inactivation. If, however, the vesicle protein usage controls the inactivation, then chronically increased synaptic activity should lead to a faster inactivation. We tested therefore the fraction of the vesicles that were still recycling after incubation for 12 hours with the GABA_A_ receptor antagonist bicuculline, or with a Ca^2+^ concentration raised to 8 mM. Both treatments lead to a chronic increase in synaptic activity (~2-fold, see Supplemental Experimental Procedures). To test their effects, we first incubated the neurons for 1 hour with unconjugated synaptotagmin 1 antibodies, to tag the entire recycling pool. We then applied the drugs, and after 12 hours applied Cy5-conjugated secondary antibodies onto the living neurons to detect the still recycling molecules, followed by fixation, permeabilization, and application of Cy3-conjugated antibodies, to detect all other molecules (same procedure as in Supplementary Fig. 2). We found that both bicuculline and 8 mM Ca^2+^ treatments resulted in a decrease of the amount of molecules that still participated in recycling, coupled to an increase of the amount of inactivated molecules. This implies the existence of a mechanism that acts as a timer for synaptic vesicle release, and inactivates the “used” vesicles after a certain amount of release.

The data we gathered so far allowed us to model the vesicle protein life cycle mathematically (Supplementary Fig. 7). As assumptions for this model, we used the result presented above, that newly synthesized synaptic vesicle proteins start out in the releasable population. The synaptic vesicle protein assemblies retrieved upon recycling can then become inactivated, a state from which they would enter the degradation pathway (Supplementary Fig. 7a), without returning to the recycling state. The model we constructed from this is based on a series of exponential equations (see Online Methods), and recapitulates the data we presented so far on synaptic vesicle degradation (Supplementary Fig. 7b). The model thus enabled us to predict the probability distributions of the time it takes to inactivate synaptic vesicle proteins (Supplementary Fig. 7c), the time the inactivated vesicles remain in the inactive population within the synapse before degradation (Supplementary Fig. 7d), the total synaptic vesicle protein lifetime (Supplementary Fig. 7e), and the usage of synaptic vesicle proteins during their lifetime (Supplementary Fig. 7f). The average number of release rounds per synaptic vesicle protein lifetime predicted from the model is ~199.

This number can also be cross-validated with values obtained by a different approach, based on measuring three essential parameters: the frequency of synaptic activity in culture during undisturbed network activity, the percentage of the vesicle proteins from the active pool that recycle during each synaptic activity burst, and the amount of time spent by the vesicle proteins in the releasable population. We had already obtained the last parameter from Supplementary Fig. 3c. The neuronal activity rate was measured by monitoring neuronal activity bursts, using the calcium indicator construct GCaMP6 (Chen et al., 2013) (Fig. 5a-c). The frequency of the bursts of activity was ~0.09 Hz (Fig. 5c). To estimate the percentage of the vesicle proteins released per activity burst, we performed simultaneous imaging of GCaMP6 and the synaptophysin-based pHluorin sypHy (Granseth et al., 2006; Li et al., 2011), as an indicator of synaptic vesicle release (Fig. 5e-g). Synaptic vesicle release robustly coincided with Ca^2+^ bursts (Fig. 5f,g). Each burst triggered the release of ~2-3% of all sypHy molecules, on average (Fig. 5h,i). From the activity burst frequency, the protein fraction released per activity burst, and the time constant of inactivation (Supplementary Fig. 3), one can calculate the average number of release events that a set of vesicle molecules undergoes under intrinsic network activity conditions. This averages to ~210 rounds of release (see Online Methods for the calculation), which is very close to the result obtained with the considerably less biased model from Supplementary Fig. 7.

**Figure 5:**
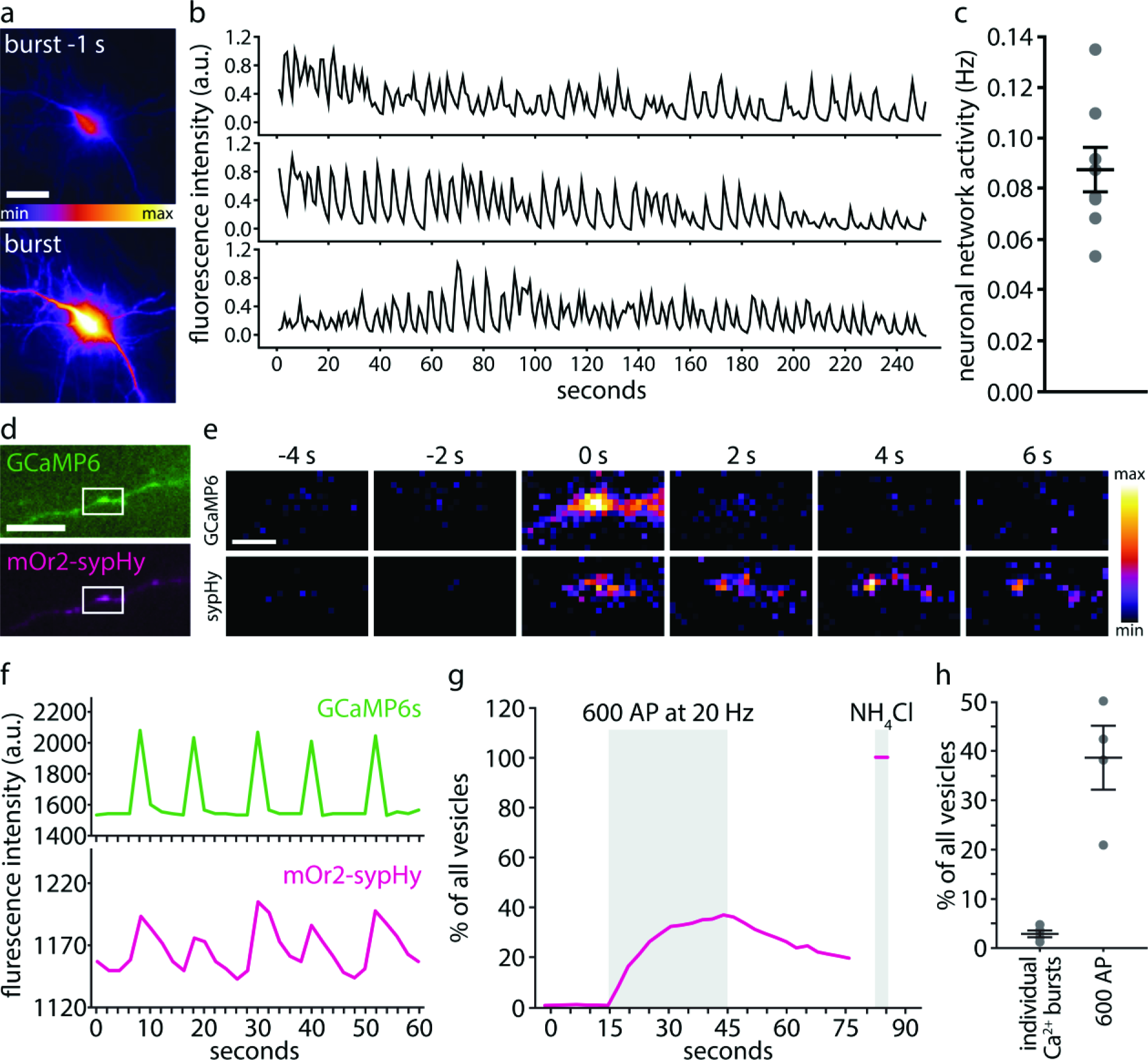
An analysis of the synaptic vesicle recycling under intrinsic network activity. (a) To determine the frequency of neuronal activity under intrinsic (non-stimulated) conditions, neurons were transfected with the calcium sensor GCaMP6 for 3-4 days, and then imaged using a Nikon Ti-E microscope (epifluorescence). The panels show typical images of a neuron during a high activity phase (burst; high Ca^2+^ concentration) or during a low activity phase (burst −1 s; low Ca2+ concentration). Scale bar 20 μm. (b) Exemplary traces of intrinsic network activity in hippocampal cultures. The fluorescence signal from cells such as those in panel (a) was measured for several minutes, and is shown here as arbitrary fluorescence units. (c) We analyzed the average frequency of the activity bursts in hippocampal cultures (n = 7 experiments), from traces as those shown in panel (b). (d) To determine the fraction of the vesicles that recycle during one burst of activity, we co-transfected neurons with GCaMP6 and with the vesicle recycling sensor sypHy, and we then assessed the increases in sypHy fluorescence coinciding with activity bursts. Scale bar 2 μm. (e) Exemplary images of changes in GCaMP6 and sypHy fluorescence during a pulse of intrinsic network activity (white box magnified from (e)). (f) Exemplary traces of activity (GCaMP6 signal) and exocytosis levels (sypHy signal) in hippocampal cultures. (g) As a comparison to the spontaneous exocytosis measurements from (d-f), we show here an exemplary trace indicating exocytosis (sypHy signal) during a train of electrical stimulation with 600 action potentials (AP) at 20 Hz. This stimulus releases the entire recycling pool (Wilhelm et al., 2010). To assess the fraction of the vesicles that were released (the fraction of the sypHy molecules that are located in the recycling pool), a pulse of NH_4_Cl was applied at the end of the experiment, which reveals all sypHy molecules, both in the recycling and in the inactive pool. (h) We quantified the amount of sypHy exocytosed during one activity burst, from traces as shown in panel (g), as percentage of the total amount of sypHy, revealed by a pulse of NH_4_Cl, as in panel (h). As a comparison, the recycling pool, revealed by stimulation with 600 action potentials (AP) at 20 Hz (as in panel h) is shown. (n = 4 experiments for both conditions).

### Inactivation may be triggered by contamination with SNAP25

A timer mechanism that can estimate the number of release events could be based on changes in the composition of the vesicles. For example, small amounts of proteins could be eventually lost or gained from the vesicle protein assemblies during recycling (see Supplementary Fig. 1 and the introductory paragraphs), thereby resulting in changes in the composition of the recycling vesicles.

To test for this, we labeled the recycling pool of vesicles by incubating the neurons with Atto647N-conjugated synaptotagmin 1 antibodies. We then either fixed immediately the neurons, thus maintaining the fixed synaptotagmin 1 molecules in the recycling pool, or incubated the neurons for a further 4 days, to ensure that the molecules switched to the inactive pool. We then immunostained the cultures for several candidate proteins, and imaged them by 2-colour STED microscopy, using a Leica TCS STED microscope. To ensure excellent Z-resolution, the cultures were embedded in melamin resin, and were cut into in ultrathin sections before imaging, as we performed in the past for imaging such preparations (Punge et al., 2008). Finally, we analyzed the co-localization of individual synaptotagmin 1 spots with the different markers.

We chose the candidate proteins based on their abundance (Takamori et al., 2006; Wilhelm et al., 2014), based on their importance in synaptic vesicle release (Jahn and Fasshauer, 2012; Rizzoli, 2014), and based on their presence in compartments involved in synaptic vesicle release and recycling (Rizzoli, 2014; Wilhelm et al., 2014). The following candidate proteins were tested: SNAP25 and syntaxin 1 for the cell membrane, VGlut 1/2, vATPase, VAMP2 and synaptotagmin 1 for the synaptic vesicles themselves, syntaxin 16 and VAMP4 for endosomal compartments, and synapsin as an abundant soluble vesicle-associated protein. We detected only one significant change: the co-localization of the synaptotagmin 1 spots with SNAP25 was higher for the ageing synaptic vesicles (Fig. 6a), which had approximately 2-fold more SNAP25 than young ones. This presumably takes place via SNAP25 being picked up during vesicle retrieval from the plasma membrane, where SNAP25 is present at a 6-7 fold higher density (per μm^2^ of membrane) than in synaptic vesicles (Takamori et al., 2006; Wilhelm et al., 2014). Interestingly, its functional partner, syntaxin 1, is present at almost equal density in synaptic vesicles and in the plasma membrane (Takamori et al., 2006; Wilhelm et al., 2014), suggesting that there is virtually no possibility of contaminating vesicles with syntaxin 1, but that this is possible for SNAP25.

**Figure 6:**
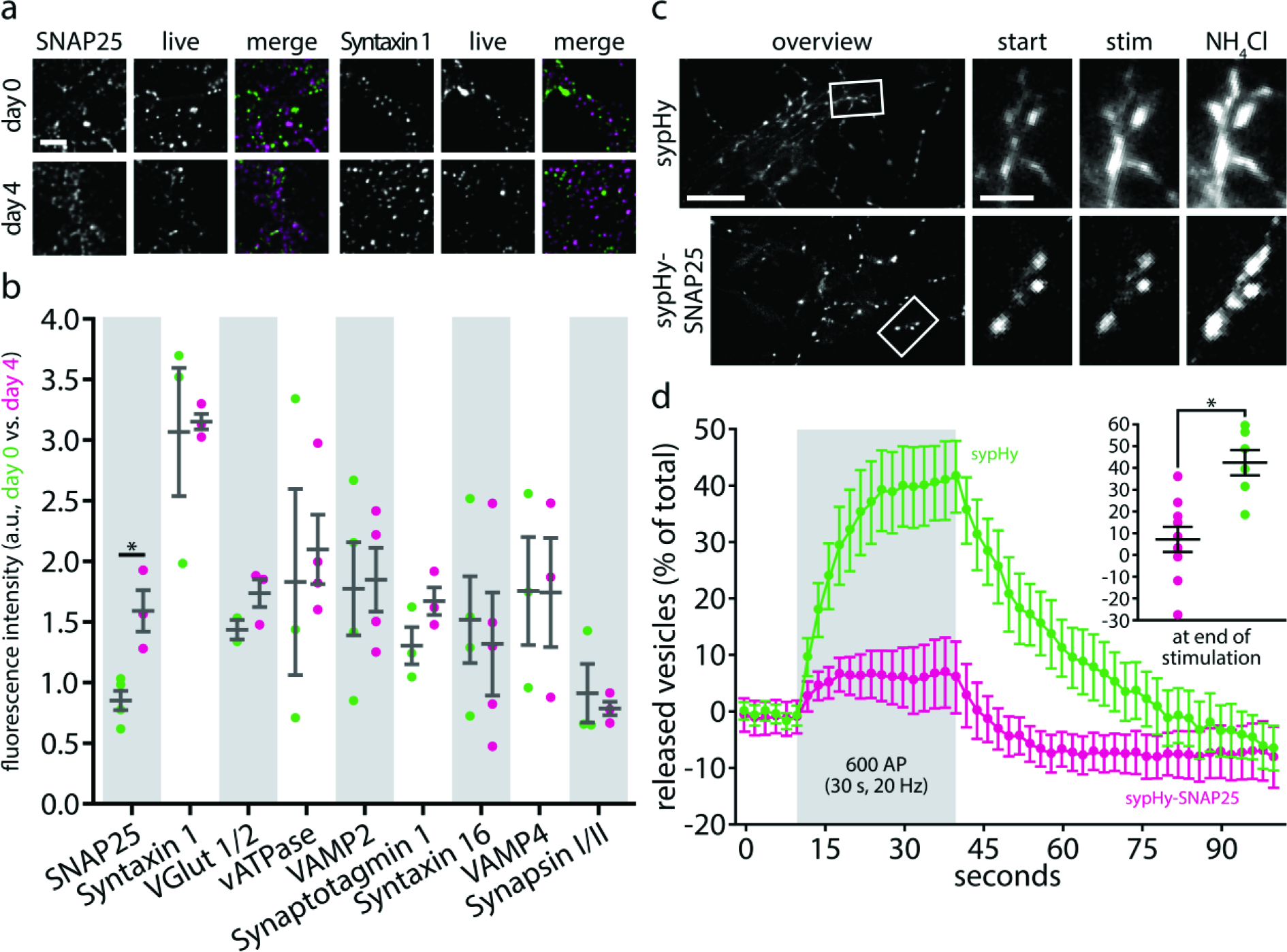
SNAP25 enters ageing synaptic vesicles and inactivates them. (a-b) Two-color STED analysis of changes in synaptic vesicle protein levels during the transition from the releasable state to the inactive state. Living neurons were incubated with Atto647N-conjugated synaptotagmin 1 lumenal domain antibodies for 1 hour at 37°C, to label the actively recycling vesicles, as in Figure 1a, and were then fixed and co-immunostained for different proteins of interest directly, or were placed in a cell culture incubator for 3-4 days, to enable the antibody-labeled molecules to enter the inactive pool, before fixation and co-immunostaining. The samples were embedded in melamin and cut in ultrathin (50 nm) sections, before 2-color STED imaging, as in Figure 2b. Exemplary images are shown in (a) for syntaxin 1 and SNAP25, with the protein of interest signal next to the signal from the live-tagging of synaptotagmin 1, and a merged image of both, for day 0 (releasable vesicles) and day 4 (inactive vesicles); scale bar 1 μm. We selected abundant and functionally important proteins of interest from every compartment the vesicle passes through during recycling (synaptic vesicle, endosome, cell membrane). We analyzed the amount of fluorescence corresponding to the protein of interest that overlapped with the synaptotagmin 1 signal (i.e. the two signals were presented within the same voxels, which are substantially below the synaptic vesicle volume in this experiment) in (b). The only protein whose levels changed significantly is SNAP25 (SNAP25, n (day 0) = 4, n (day 4) = 3, *p = 0.0124, t(5) = 3.8200; Syntaxin 1, n (day 0) = 3, n (day 4) = 3, p = 0.8850, t(4) = 0.1541; VGlut 1/2, n (day 0) = 2, n (day 4) = 3, p = 0.1986, t(3) = 1.6447; vATPase, n (day 0) = 3, n (day 4) = 4, p = 0.7340, t(5) = 0.3594; VAMP2, n (day 0) = 4, n (day 4) = 4, p = 0.8837, t(6) = 0.1527; Synaptotagmin 1, n (day 0) = 3, n (day 4) = 3, p = 0.1604, t(4) = 1.7208; Syntaxin 16, n (day 0) = 4, n (day 4) = 4, p = 0.7406, t(6) = 0.3468; VAMP4, n (day 0) = 3, n (day 4) = 3, p = 0.9863, t(4) = 0.0183; Synapsin I/II, n (day 0) = 3, n (day 4) = 3, p = 0.6638, t(4) = 0.4685; at least 10 neurons sampled per experiment). (c-d) To test whether SNAP25 is able to inhibit synaptic vesicle recycling, we expressed in neurons either sypHy, or a sypHy-SNAP25 construct that targets SNAP25 to synaptic vesicles. The cells were then stimulated with 600 action potentials at 20 Hz, to trigger the release of the entire recycling pool, at 3-4 days after transfection. Exemplary images are shown in (c) for both constructs, with overviews before stimulation and magnified areas before stimulation (start), at the peak of stimulation (stim), and after application of NH_4_Cl; scale bars 20 μm (overviews) and 5 μm (magnified areas). NH_4_Cl was used to reveal all sypHy moieties, to be able to represent the exocytosis response as % of all sypHy molecules (d). The cells expressing sypHy-SNAP25 displayed a strongly reduced response (n = 6 independent experiments for sypHy, n = 9 independent experiments for sypHy-SNAP25; *p = 0.0025, t(13) = 3.7334). All imaging was performed at 37°C, with a Nikon Ti-E microscope (epifluorescence). All data represent the mean ± SEM.

This experiment suggests that SNAP25 contamination may separate young from aged vesicle protein assemblies or vesicles. This, however, does not directly imply that SNAP25 has any functional relevance on the aged vesicles. To test this specifically, we engineered a construct to target SNAP25 to synaptic vesicles: sypHy-SNAP25, a simple fusion of the synaptophysin-based pHluorin sypHy and SNAP25. Synaptophysin targets to synaptic vesicles more reliably than most other proteins (Rizzoli et al., 2006; Takamori et al., 2006), and can therefore efficiently incorporate SNAP25 into the vesicles. At the same time, the pHluorin moiety enabled us to directly observe the response of these tagged vesicles to stimulation. Exocytosis was severely suppressed by the addition of SNAP25 on the vesicles (Fig. 6b). This suggests that the SNAP25 contamination on aged synaptic vesicles is sufficient to inactivate them.

### The SNAP25 effects can be removed by expressing CSPα

The fashion in which SNAP25 could participate in inactivating vesicles is not immediately obvious. The best-known SNAP25 interactor on the vesicle is VAMP2. SNAP25 on the cell membrane interacts in *trans*-complexes with VAMP2 on the vesicle during exocytosis, so it could, in principle, be envisioned that SNAP25 on the vesicles might block all copies of VAMP2 on the vesicle in *cis*-SNARE complexes. However, VAMP2 is present in ~70 copies per vesicle (Takamori et al., 2006; Wilhelm et al., 2014), while we estimate a maximum of ~5 SNAP25 copies on the aged vesicles, assuming that vesicles start out with 1-2 copies of SNAP25 (Takamori et al., 2006), an amount which then doubles during ageing (Fig. 6a). Five SNAP25 copies are unlikely to result in sufficient interference with 70 copies of VAMP2.

However, SNAP25 is also known to interact with CSPα, a chaperone needed in the priming process, to prepare the fusogenic *trans*-complex of SNAP25 and VAMP2 (Evans et al., 2003; Jahn and Fasshauer, 2012). There are only 2-3 copies of CSPα on each vesicle (Takamori et al., 2006; Wilhelm et al., 2014). This value is far closer to our estimated number of SNAP25 copies in aged vesicles, and it is therefore conceivable that SNAP25 might sequester all of them in non-functional *cis*-complexes on the vesicle surface. Normally, a *trans*-complex forms between vesicular CSPα, SNAP25 from the plasma membrane, and two soluble molecules, the ubiquitous chaperone Hsc70 and SGT? (Sharma et al., 2012). This *trans-complex* is involved in priming SNAP25 for fusion. The formation of this complex in *cis*, on the vesicle surface, could create a quantitative bottleneck for the fusion of the aged vesicles in the form of sequestration of CSPα. According to this hypothesis, overexpressing CSPα would remove this bottleneck, and would thus remove the timing mechanism that inactivates ageing synaptic vesicles.

To test this hypothesis, we tested the size of the total recycling pool, as obtained from incubating neurons with Atto647N-conjugated antibodies for 1 hour, in different conditions. Overexpression of CSPα resulted in a substantial increase of the recycling pool, almost to the level of total pool of molecules at the synapse (Fig. 7a,b). Thus, CSPα seems indeed to remove the bottleneck mechanism that inactivates vesicles. We did not observe any change in activity levels when overexpressing a mutated form of CSPα that does not target to vesicles correctly (Sharma et al., 2011) (Fig. 7b).

**Figure 7:**
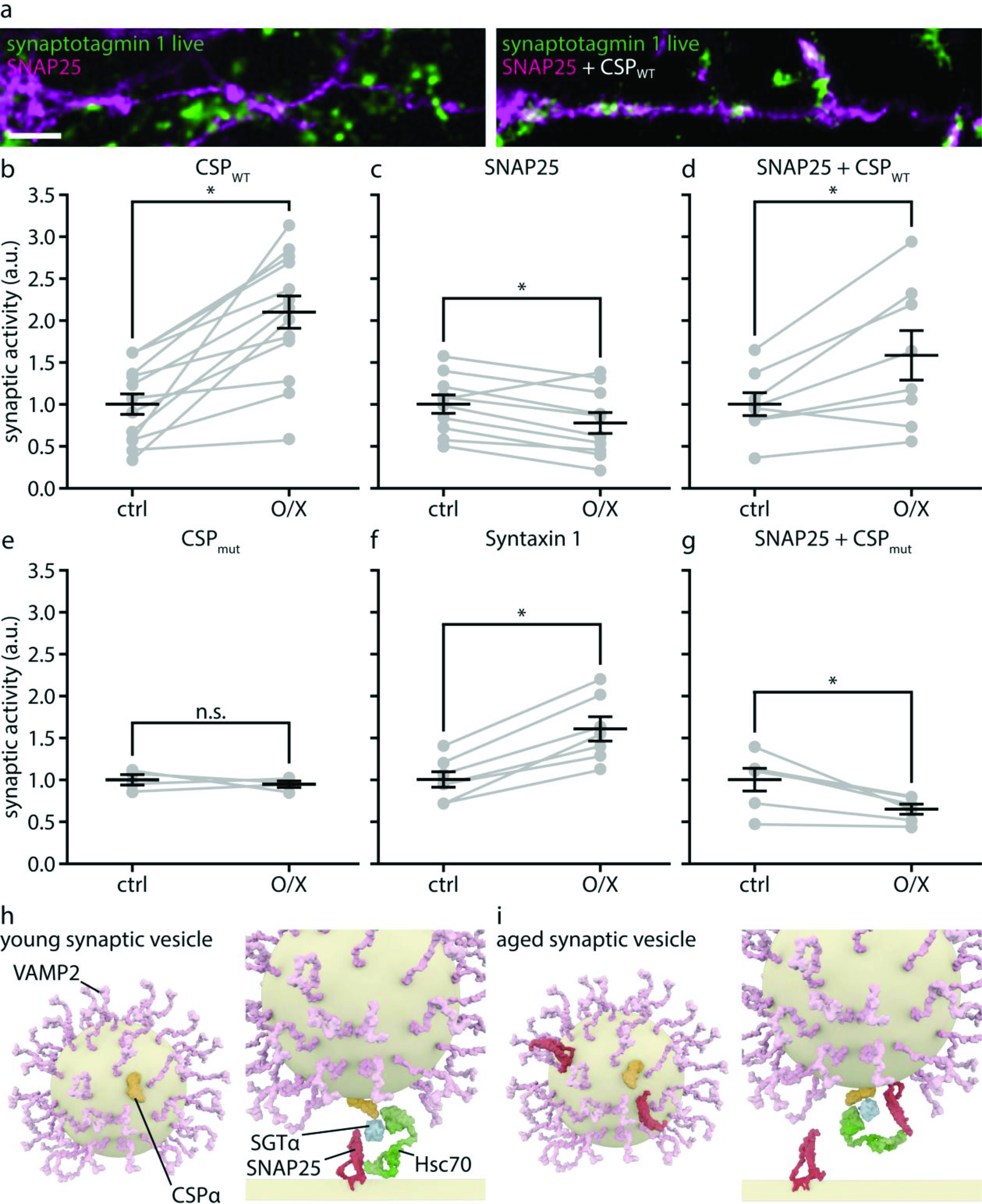
SNAP25 inactivates synaptic vesicles by blocking CSPα on the vesicle. (a) Neurons were transfected with wild-type SNAP25 (left panel) or co-transfected with SNAP25 and with wild-type CSPα (CSP_WT_, right panel). The transfected cells are shown in magenta, which corresponds to a YFP moiety that is coupled to SNAP25, for detection purposes. At 3-4 days after transfection the neurons were incubated with Atto647N-conjugated synaptotagmin 1 antibodies, for 1 hour, to label the actively recycling vesicles, as in Figure 1a. The neurons were then fixed, and the synaptotagmin antibody intensity from control neurites and from neurites containing the expressed proteins was analyzed. Exemplary images show reduced vesicle recycling (reduced synaptotagmin 1 antibody levels) in neurons overexpressing SNAP25 (left panel; very limited green intensity within the magenta areas), compared to neurons overexpressing both SNAP25 and CSP_WT_ (right panel). Scale bar: 2 μm. (b-g) We quantified the size of the actively recycling pool, determined by incubation with Atto647N-conjugated synaptotagmin 1 antibodies, as in Figure 1a, in neurons expressing different constructs, or combinations of constructs. Displayed are the values from neurons overexpressing the respective construct(s) (O/X) compared to the values from control neurons in the same samples that did not overexpress the construct(s) (ctrl). The following conditions were used: (b) neurons overexpressing CSPWT (n = 16 transfected neurons from 3 independent experiments); (c) SNAP25 (n = 24 transfected neurons from 8 independent experiments); (d) SNAP 25 + CSP_WT_ (n = 20 transfected neurons from 8 independent experiments); (e) CSP_mut_, a mutated version of CSPα unable to target to vesicles and thus incapable of interacting with SNAP25 (Sharma et al., 2012) (n = 11 transfected neurons from 4 independent experiments); (f) syntaxin 1, as a control (n = 13 transfected neurons from 7 independent experiments); (g) SNAP25 + CSP_mut_ (n = 20 transfected neurons from 6 independent experiments). The significance levels determined are: (b) *p = 0.0478, t(4) = 2.8201; (c) *p = 0.0139, t(14) = 2.8091; (d) *p = 0.0269, t(12) = 2.516; (e) ^ns^p = 0.5916, t(6) = 0.5664; (f) *p = 0.0378, t(12) = 2.3346; (g). *p = 0.0126, t(10) = 3.0354. All data represent the mean ± SEM. All imaging was performed with a Leica confocal microscope. (h) A hypothetical model of SNAP25 and CSPα activity. They interact with SGT? and Hsc70 in *trans*, promoting synaptic vesicle fusion (Sharma et al., 2012). (i) A hypothetical model of how SNAP25 on the aged vesicles may interact in *cis* with CSPα, thus sequestering it from its *trans* interaction with SNAP25 molecules from the plasma membrane.

At the same time, our hypothesis posited that the opposite effect, a decrease the recycling pool size, should be observed when overexpressing SNAP25, not linked to vesicles as in Fig. 6, but in the wild-type form (Fig. 7a,b). This effect, which is in agreement with previous findings on SNAP25 overexpression (Owe-Larsson et al., 1999), was indeed observed, and was specific to SNAP25, since expressing the other plasma membrane SNARE, syntaxin 1, resulted in a strong increase of synaptic recycling, which is actually the expected result for an experiment in which the copy numbers of a fusion protein are raised (Fig. 7b). Finally, CSPα overexpression also counteracted the effects of the SNAP25 overexpression, as predicted by our hypothesis (Fig. 7a,b).

The elimination of vesicle ageing by CSPα overexpression, however, was deleterious to the cells, suggesting that the inactivation of old vesicles is physiologically relevant. The cells overexpressing CSP_WT_ had a significantly higher proportion of damaged neurites than those expressing CSP_mut_ that does not target to vesicles (Supplementary Fig. 5a,b), as observed by investigating neurite morphology in neurons expressing cytosolic GFP, to be able to detect the neurites optimally.

At the same time, it appears that the endocytosis of aged vesicles is poorer than that of young vesicles (Supplementary Fig. 5c,d). Neurons treated with inhibitors of protein production or transport, as in Fig. 3, and which therefore can only recycle aged vesicles, were stimulated strongly (600 action potentials at 20 Hz), and were investigated by immunostaining for both synaptotagmin 1 and synaptophysin. The co-localization of the two molecules, as analyzed by confocal microscopy, was poorer in these neurons than in control neurons, suggesting that endocytosis and the maintenance of vesicle identity are poorer (Supplementary Fig. 5c,d). These effects mirror those found in mice lacking CSPα (Rozas et al., 2012), which suffer from neurodegeneration, and have endocytosis defects that were difficult to reconcile solely with a role of CSPα in SNAP25 priming. Our results explain this latter finding, since old vesicles would continue to recycle in the absence of CSPα, leading to defects in vesicle recycling.

### The effects of SNAP25 overexpression on vesicle degradation

The inactivated synaptic vesicles must presumably be degraded at a later time point. We hypothesized that this may occur through the direct participation of SNAP25. Synaptic vesicle degradation is widely assumed to entail fusion to the endo-lysosomal system (Katzmann et al., 2002; Raiborg and Stenmark, 2009; Rizzoli, 2014). This fusion event requires a combination of R, Q_a_, Q_b_, and Q_c_ SNAREs on the surface of the endosome and of the synaptic vesicle. While vesicular R-SNAREs (VAMP2) and endosomal Q_a_ SNAREs are present in abundance in the synapse, Q_b_- and Q_c_-SNAREs are relatively scarce (Wilhelm et al., 2014). SNAP25 is a Q_bc_ SNARE, and might thus increase the probability of fusing the vesicles to the endo-lysosomal system, as has been previously suggested in PC12 cells (Aikawa et al., 2006a, 2006b). We therefore tested whether the overexpression of SNAP25 or sypHy-SNAP25 increases the co-localization of the vesicles with the recycling endosome marker Rab 7, which is thought to be the first step in vesicle degradation. This was observed in both cases by verifying the co-localization of Rab 7 with the synaptic vesicle marker synaptophysin, in conventional immunostaining experiments (Supplementary Fig. 6).

## DISCUSSION

Organelle ageing has long been recognized as an important factor in cellular disease and death. However, little is known about how aged organelles are identified before they can noticeably impair cellular pathways, especially for post-mitotic cells, which are most strongly affected by malfunctioning organelles. We address here this problem for a well understood organelle, the synaptic vesicle. There is substantial information on how the synaptic vesicle is degraded (Binotti et al., 2014; Fernandes et al., 2014; Rizzoli, 2014; Uytterhoeven et al., 2011), but it has never been clear why a fraction of the vesicles are inactivated and are reluctant to participate in neurotransmission (Alabi and Tsien, 2012; Denker and Rizzoli, 2010; Denker et al., 2011a; Rizzoli, 2014; Rizzoli and Betz, 2005). We suggest here that this mechanism is the gradual contamination of synaptic vesicles from the recycling (active) pool with SNAP25, during multiple rounds of release and recycling.

This mechanism fits most easily with the concept that the identity of the vesicle is maintained for a relatively long time, throughout multiple rounds of recycling, as discussed in the introduction. However, it also fits with the view of intermixing of vesicle components in the plasma membrane, after exocytosis. In this view, meta-stable vesicle protein assemblies, albeit not full vesicles, can become contaminated with SNAP25 upon recycling. The contaminated assemblies are then incorporated into full vesicles during endocytosis, and the resulting vesicles are thereby tagged as old vesicles, become inactive, and are eventually degraded.

### The synaptic vesicle life cycle

Based on our data, we suggest the following sequence of events: synaptic vesicle precursors are produced in the soma, are transported to the synapse, where they are assembled into releasable vesicles. The vesicle proteins are used in exocytosis for up to a few hundred times during their lifecycle (Fig. 7c), and the protein assemblies get inactivated by contamination with SNAP25, which blocks CSPα in futile *cis*-complexes (Fig. 7d), before the vesicles are ultimately degraded.

Our results offer a new interpretation to the long-standing discussion on molecular differences between releasable and inactive “reserve” vesicles (Rizzoli and Betz, 2005). The difference between the two pools seems to be the age and usage of the vesicle molecules, measured by the accumulation of SNAP25 within their assemblies. Vesicles that work numerous times are identified by this mechanism, and are removed from the recycling pathway before they can become so badly damaged as to endanger the function of the synapse, and the organism’s survival along with it. The inactive vesicles, which are reluctant to release and can only be forced to exocytose under strong supra-physiological stimulation, may not act as a reserve for neurotransmission, but are a collection of aged vesicles that are prevented from releasing. These vesicles may take on different roles in their late age, such as providing a buffer capacity for soluble co-factors of synaptic vesicle exo-and endocytosis (Denker et al., 2011b).

### SNAP25 and CSPα as molecular timer for synaptic vesicle inactivation

The role of SNAP25 in the SNARE complex that facilitates synaptic vesicle fusion to the cell membrane is well established (Jahn and Fasshauer, 2012), and CSPα has long been studied as a major chaperone involved in promoting the formation of the SNARE complex (Evans et al., 2003; Sharma et al., 2011, 2012). The knockout of SNAP25 leads to a complete failure of neurotransmission and death (Washbourne et al., 2002), as expected for such a critically important protein. The knockout of CSPα has much milder effects, albeit the mice develop neurological problems that lead to death within 1-2 months (Fernández-Chacón et al., 2004).

CSPα has been suggested to be mainly needed in priming SNAP25 for exocytosis, and presumably also in folding and stabilizing this protein. Without CSPα, exocytosis upon single action potential stimulation is somewhat poorer, as would be expected from less efficient priming (Rozas et al., 2012; Sharma et al., 2012). This phenotype is accompanied by a loss of SNAP25, since the stabilization provided by CSPα is eliminated. Along the same lines, the overexpression of CSPα has been shown to help stabilize SNAP25, and to result in more synaptic vesicle fusion (Sharma et al., 2011), as in our experiments (Fig. 7).

More unexpectedly, endocytosis is also poorer in mice lacking CSPα (Rozas et al., 2012), which was difficult to ascribe to a role of CSPα in priming or stabilizing SNAP25. Our results provide a simple interpretation to this finding: older vesicles are more poorly retrieved (Supplementary Fig. 5c,d), which necessarily results in an endocytosis defect.

At the same time, these findings suggest that old vesicles are not inherently unable to release. They are only less able to do so than young vesicles, since they do not prime as efficiently as the young ones. When a young vesicle approaches the active zone, SNAP25 from the plasma membrane interacts with the CSPα from the vesicle surface, and is primed and readied for fusion. In contrast, for an old vesicle the CSPα molecules are less likely to prime the plasma membrane SNAP25, since they can alternatively interact with the vesicle-bound SNAP25. This makes such vesicles less able to prime, and probably prevents them from docking, as long as young vesicles are abundant in the vicinity. Especially under conditions of strong stimulation, however, where the young vesicles are all rapidly depleted, the aged vesicles could still be recruited to release, albeit with ever decreasing efficiency (Fig. 1d).

### The inactivation of synaptic vesicles precedes the accumulation of damage to their proteins

One caveat of the work presented above is that, while there is evidence that SNAP25 is able to inactivate the vesicles, it cannot be concluded that this is the only such marker on an aged vesicle. It is possible that other elements play a role as well, such as the accumulation of oxidative damage. However, it is currently not possivle to directly measure the damage, such as oxidation, suffered by individual molecules on individual vesicles, in a live-cell experiment.

To obtain more insight into this issue, we turned to previous investigations of vesicle protein lifetimes. Currently, there is little evidence available on how fast synaptic proteins might accumulate damage. However, if we assume that degradation of proteins occurs only after they have been damaged, the lifetimes of synaptic proteins in culture (Cohen et al., 2013) can be used to predict the rates of damage to synaptic vesicle proteins (see Supplemental Experimental Procedures). We plotted the cumulative prediction of synaptic vesicle protein damage, and explored how much of the protein complement of one synaptic vesicle would be damaged at the time of inactivation (Supplementary Fig. 8). This calculation predicts that virtually no synaptic vesicle proteins are damaged at this time point, and that synaptic vesicles are removed just before the accumulation of damage begins. This indicates that the molecular timer we identified probably acts as a predictive mechanism, which pre-emptively removes vesicles from neurotransmission, before they can be damaged and become a hazard to cellular function, as outlined in the Introduction.

It is still unclear whether all membrane proteins of the aged vesicle will be degraded simultaneously in the cell body, or whether some, which are not yet damaged, will escape degradation. Such proteins could be again used in the formation of synaptic vesicles, as has been suggested in the past for dense-core vesicles (Vo et al., 2004), but this issue requires further investigation.

### Conclusion

We conclude that the synapse evolved to accurately predict and pre-empt synaptic vesicle damage, using a molecular timer involving SNAP25 and CSPα. Synaptic vesicles are removed from activity just as they are expected to start accumulating damage. This is necessary since the cell depends on a fairly small population of releasable synaptic vesicles, which should not be compromised by even the slightest damage, if they are to ensure continued and reliable neurotransmission. We suggest that such mechanisms might be present in other cellular processes as well, especially if they depend on the action of only a few organelles at a time.

## METHODS

Methods, including statements of data availability and any associated accession codes and references, are available in the Supplementary Information.

**Figure 4:**
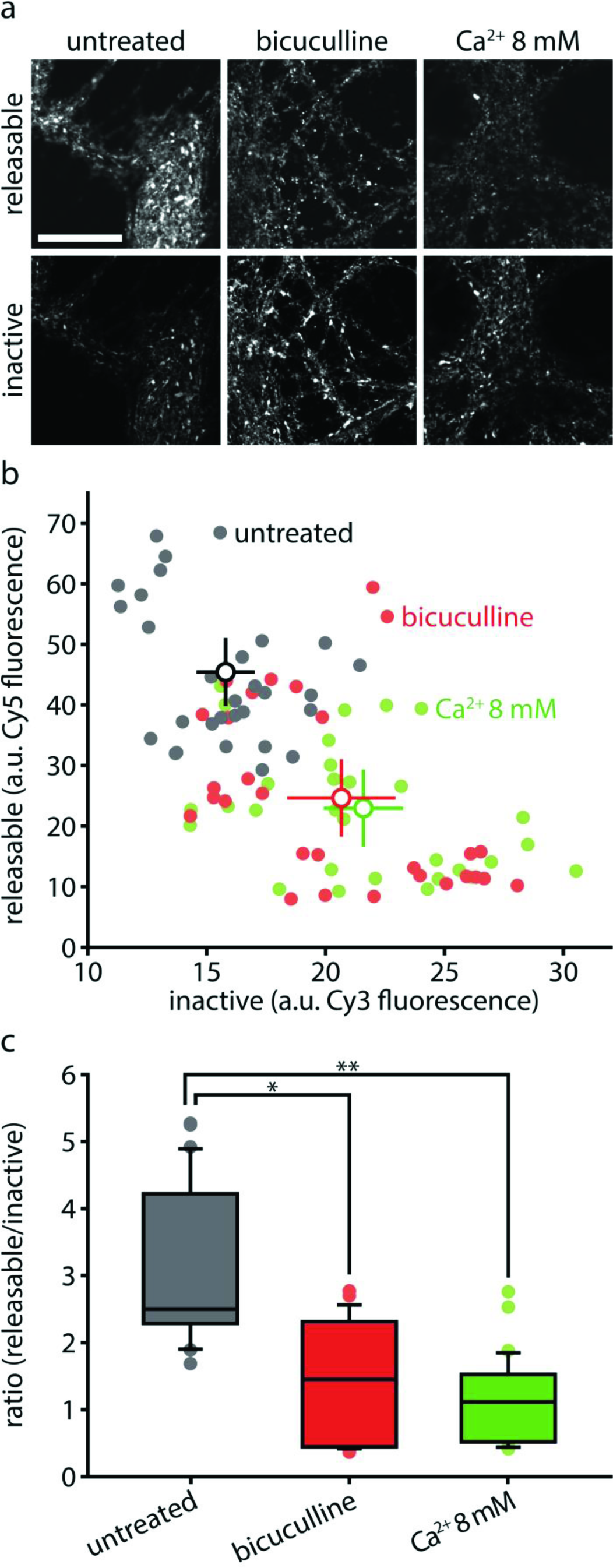
Increased synaptic activity accelerates synaptic vesicle ageing. (a-b) We investigated the effect of increasing neuronal activity on the ageing and inactivation of vesicles as follows. We tagged synaptotagmin 1 epitopes with non-conjugated antibodies, and then, after a 12 hour interval, we applied fluorophore-conjugated secondary antibodies (Cy5) to the living cultures for 1 hour. These antibodies revealed all of the synaptotagmin 1 antibodies that were exposed during vesicle recycling. We then fixed and permeabilized the neurons, and applied secondary antibodies conjugated to a different fluorophore (Cy3), to thereby reveal all of the remaining, no longer releasable, synaptotagmin 1 antibodies. This procedure thus indicates the proportion of the tagged synaptotagmin 1 epitopes that are still recycling in the synapses under normal activity, as well as the proportion that are present in the synapses. Exemplary images of releasable and inactive vesicles are shown in (a); scale bar 20 μm. Data from images such as shown in (a) was quantified, with the fluorescence intensity from releasable (Cy5) and inactive (Cy3) epitopes plotted in (b). The experiment was performed in untreated control cultures, or in cultures incubated for the 12 hours with bicuculline or 8 mM Ca^2+^ to increase synaptic activity (n = 30 neurons for control, 29 neurons for bicuculline, 30 neurons for Ca^2+^ 8mM, from 3 independent experiments per condition). The intensity of the signal ascribed to releasable or inactive vesicles is shown. (c) Ratio of releasable vs. inactive vesicles, in arbitrary units, from (a); *p = 0.0001, t(57) = 6.2692; **p = 0.0001, t(58) = 7.8419. All data represent the mean ± SEM. Imaging was performed with a Leica confocal microscope.

## ACKNOWLEDGEMENTS

We thank Reinhard Jahn for providing a plasmid for YFP-SNAP25. We thank Erwin Neher for help with the development of the mathematical model of the synaptic vesicle life cycle. We thank Martin Meschkat, Andreas Höbartner, Annedore Punge, and Peer Hoopmann for help with the experiments. We thank Burkhard Rammner for providing the illustrations of synaptic vesicle and protein dynamics. We thank Manuel Maidorn, Martin Helm, and Katharina N. Richter for critically reading the manuscript. S.T. was supported by an Excellence Stipend of the Göttingen Graduate School for Neurosciences, Biophysics, and Molecular Biosciences (GGNB). E.F.F. is a recipient of long-term fellowships from the European Molecular Biology Organization (ALTF_797-2012) and from the Human Frontier Science Program (HFSP_LT000830/2013). The work was supported by grants to S.O.R. from the European Research Council (ERC-2013-CoG NeuroMolAnatomy) and from the Deutsche Forschungsgemeinschaft (Cluster of Excellence Nanoscale Microscopy and Molecular Physiology of the Brain, as well as DFG RI 1967 2/1). The nanoSIMS instrument was funded by the German Federal Ministry of Education and Research (03F0626A).

## AUTHOR CONTRIBUTIONS

S.O.R., S.T., and A.D. conceived the project. S.J. and A.V. (Leibniz Institute) performed the nanoSIMS experiments. E.F.F. performed the experiments on culture activity with GCaMP6 and sypHy. A.V. (Cells in Motion Cluster) performed the two-color STED experiments on changes in synaptic vesicle protein levels. S.T. performed all other experiments. H.W. developed the mathematical model of the synaptic vesicle life cycle. S.O.R., S.T., and E.F.F. analyzed the data. S.O.R. and S.T. prepared the manuscript.

